# “Mapping of Gonadal Development in Cryptorchidism: UTF-1 and Germ Cell Dysgenesis”

**DOI:** 10.64898/2026.06.24.734274

**Authors:** Pablo A. Suarez, Mariel Magdits, Mei Cao, Cornelia Ding, James F. Smith, Laurence S. Baskin, Yi Li

## Abstract

**Study question:** How does cryptorchidism affect germ cell development and UTF-1–mediated pluripotency potential at the time of orchiopexy?

**Summary answer:** Cryptorchidism was associated with the following germ cell abnormalities: germ cell clustering with many cords/tubules lacking germ cells and reduced UTF-1 expression, suggesting limited germ cell differentiation into spermatogonia

**What is known already:** Cryptorchidism, affecting 1.6-9% of male newborns, is known to increase the risk of infertility and testicular cancer due to abnormal germ cell development. Germ cells and pluripotent stem cell gene, UTF-1, play critical roles in spermatogonia differentiation, self-renewal, and spermatogenesis. No prior study has evaluated the testicular development by immunohistochemically mapping of these cell populations,

**Study design, size, duration:** A cross-sectional study of 31 postnatal cryptorchid testis and 5 age-matched scrotal testicular biopsies obtained from UCSF’s pathology department performed between 1993–2023.

**Participants/materials, setting, methods:** Specimens were grouped by age at surgery (6-18 months, 19 months-7 years, 8-12 years, and ≥13 years) and testis location (palpable vs. non-palpable). Scrotal prepubertal testis biopsies were sourced through the Pedi-LIFE program, a fertility preservation research biobank, with at least one control per age group. Immunohistochemistry was performed to stain specimens for germ cell (DDX4, OCT4, TSPY), pluripotent cell marker (UTF-1), as well as other key testis cell markers (A-actin, AR, P450, Sox-9), with staining graded based on control expression levels. The number of germ cells per seminiferous tubule was quantified and compared across anatomical locations using appropriate statistical analyses.

**Main results and the role of chance:** This study included 36 specimens, comprising 31 cryptorchid testes (86%) and 5 scrotal control testes (16%). The cryptorchid group exhibited testicular dysgenesis and reduced germ cell expression, correlated with increased age and testis location. Qualitative assessment revealed reduced germ cell expression across all ages in cryptorchid testes. The number of germ cells per tubule was markedly reduced in cryptorchid compared with scrotal testes after 19 months of age for DDX4, TSPY, and UTF-1. Germ cell clusters were identified in 15 out of 31 cryptorchid specimens (48%) stained for DDX4 and TSPY. UTF-1 expression was lower in cryptorchid testes across all age groups. No significant differences were noted in other testicular cell markers.

**Large scale data:** NA

**Limitations, reasons for caution:** First, the power and generalizability of the study are limited by the availability of specimens within each age group, particularly for scrotal testes, as biopsies of these tissues are not routinely performed. Second, a cross-sectional study design limits a longitudinal comparison to evaluate changes in marker expression, delayed maturation, or irreversible germ cell loss. Third, immunohistochemistry data is semi-quantitative, and protein detection is affected by antibody sensitivity and tissue preservation and influenced by antibody sensitivity. Lastly, scrotal testis used as controls were obtained from cryopreserved tissue from patients with other unrelated pathology, which may influence histological profiles.

**Wider implications of the findings:** Collectively, our findings support a model in which cryptorchidism involves both germ cell depletion and disrupted SSC lineage formation, with UTF-1 downregulation and germ cell clustering as early signatures of testicular dysgenesis. These features may help identify high-risk patients for worsening gonadal dysgenesis and infertility and can provide a rationale for earlier orchiopexy or SSC-preserving strategies.

**Study funding/competing interest(s):** The authors declare no conflicts of interest and received no funding for this study.

**Data Availability Statement:** The data underlying this article cannot be shared publicly due to ethical and legal restrictions related to the use of human tissue specimens, which may compromise donor privacy and confidentiality. Data are available from the corresponding author upon reasonable request and subject to institutional and ethical approvals.

## Introduction

Cryptorchidism, or undescended testes (UDT), is one of the most prevalent congenital anomalies, affecting 1.6-9% of male newborns^1^. Gonadal development relies on the coordination of hormonal, genetic, and environmental factors that regulate testicular descent and the establishment of spermatogenesis ^2^. A critical part of testicular development is the maturation of spermatogonial stem cells (SSCs), which are essential for maintaining spermatogenesis.^3^ This process involves precise genetic and epigenetic regulation, ensuring the proper germ cell differentiation and function ^3^. In cryptorchidism, disrupted microenvironments impair this process, leading to abnormal germ cell development, reduced fertility, and a higher malignancy risk ^4^.

Increased temperatures in undescended testes can negatively interfere with SSC differentiation and growth, potentially resulting in defective spermatogenesis and a higher risk of testicular cancer ^5^. Moreover, disturbances in the spatial and temporal organization of this environment can cause variations in germ cell characteristics and decreased viability, further affecting fertility ^6,7^. Studies indicate that the failure of germ cells to differentiate properly during mini-puberty results in diminished proliferation and delayed gonocyte clearance, contributing to long-term reproductive consequences ^4^. To better understand the molecular mechanisms underlying these developmental disruptions, it is useful to examine the expression and function of specific markers associated with germ cells, Leydig cells, Sertoli cells, and pluripotency in cryptorchid testis compared to scrotal testis.

Germ cells are known to undergo apoptosis in UDT and are associated with a proportional reduction in expression with age at the time of surgery ^8,9^. Prior work has demonstrated that DDX4 and TSPY are important genes for germ cell proliferation, spermatogonia survival and effective spermatogenesis ^10,11^. More recently, UTF-1 has been identified as a novel pluripotency marker with studies showing that its inactivation results in fewer gonocytes, defective spermatogenesis, and reduced sperm count ^12,13^. Despite its significance in early gonadal development, research on UTF-1 in the context of cryptorchidism is still limited, making it a crucial but understudied factor in testicular development and reproductive health.

Our study aims to address these gaps by providing a comprehensive immunohistochemical analysis of germ cell and pluripotency markers in both orthotopic and cryptorchid testes. By categorizing specimens based on age and testis location, we also seek to delineate how expression of these markers changes throughout development and how these changes differ between scrotal and UDT. We predict aberrant expression of spermatogonial stem cell and germ cell markers in underdeveloped cryptorchid testes compared with orthotopic testes. Our findings will provide new insights into the developmental trajectory of germ cells in the context of undescended testes and enhance our understanding of cryptorchidism.

## Methods

### Specimen Selection & Acquisition

UDT specimens were obtained from the UCSF Benioff Children’s Hospital Pathology Department from biopsies performed between 1993–2023. Pathology and surgery reports in the Electronic Health Record (EHR) were reviewed to determine age at surgery and testicular location, categorized as palpable or nonpalpable. Specimens with testicular malignancy or non-viable remnants were excluded.

Control samples (postnatal scrotal testes, n=5) were obtained from PEDI-LIFE (IRB# 13-11218), UCSF’s oncology–fertility preservation program, from patients undergoing cryopreservation due to testicular malignancy or gender dysphoria. All specimens were grouped by age at surgery and: 6–18 months (optimal surgical window based on guidelines, n=6), 19 months–7 years (pre-puberty, n=9), 8–12 years (peri-puberty, n =10) and 13+ years (post-puberty, n=16) (**Supplementary Table 1**). These ages were selected based on histological puberty age, which was estimated according to previously cited average male pubertal milestones.^14^

### Immunohistochemical Staining

For immunohistochemistry (IHC) staining, somatic, germ cell, and pluripotency markers previously mapped in the ontogeny of testis were investigated in this study; these markers were selected because their expression patterns during normal testicular development are well established in the literature.^13,15,16^ OCT-4 was selected as a negative control for our study, which has been shown to be absent at all ages of normal testis.^15^ Somatic tissue sections were deparaffinized, rehydrated through graded ethanols, and subjected to antigen retrieval using citric acid–based unmasking solution (Vector H-3300-250) with microwave treatment (25 min–1 hr). Primary antibodies were applied for 24hrs at 4°C, followed by biotinylated secondary antibodies for 1 hr at room temperature (**Supplemental Table 2**). Signal was amplified using an ABC kit (Vector PK-6100, pH 6.0) and developed with DAB (Sigma D5637). Sections were counterstained with hematoxylin, de-stained (1% HCl/EtOH), blued in ammonia water, dehydrated, cleared in Histoclear, and coverslipped with CoverSafe.

### Immunofluorescence

The same steps described for IHC above were done until the addition of the primary antibody. Primary antibodies for DDX4 (ab13840/Abcam) and UTF-1 (AF3958/ R&D Systems) were added to the sections for an incubation period of 24hrs at 4°C. Following three 5-minute washes with PBST (1× PBS containing 0.05% Tween-20) on a shaker, the sections were incubated with the appropriate secondary antibodies diluted in blocking buffer for 2hrs. After three 30-minute PBST washes, excess liquid was removed, slides were incubated in an anti-fade mounting medium containing DAPI Fluoromount-G (0100-20, SouthernBiotech, Birmingham, AL) tissue. Slides were cover slipped and sealed with nail polish.

### Specimen Imaging

Sections were photographed at 20-40x using a Leica DM 4000B microscope and Leica DFC 500 and DFC 7000T cameras using the Leica Application suite versions 4.13. The staining patterns were assessed at 20x for all images. For nuclear staining of UTF-1, a 40x magnification was used for more precise description of the results.

### Germ Cell Count

Germ cell quantification was performed at 20× magnification on testicular biopsies. For each cross-section, 10 representative seminiferous tubules were identified, and germ cells were individually counted. The primary outcome was the average number of germ cells per tubule. Counts were independently performed by two blinded observers, with discrepancies resolved by consensus.

### Ethical Approval

This study was approved by the Institutional Review Board (IRB) at the University of California, San Francisco under protocol number IRB# 19-27955. All samples used in this research were fully de-identified prior to analysis. In accordance with institutional and federal guidelines, the study qualified for exempt status and did not require informed consent due to the use of anonymized human tissue. No identifiable patient information was accessed or recorded during the duration of this study.

## Results

A total of 36 specimens were immunohistochemically stained for gonadocyte and non-gonadocyte markers and their patterns of expression were compared by age at time of surgery and testis location (**Table 1**). Of those, 5 were scrotal testis and 31 were UDT, as shown in **Supplemental Table 2**.

**Table 1.**
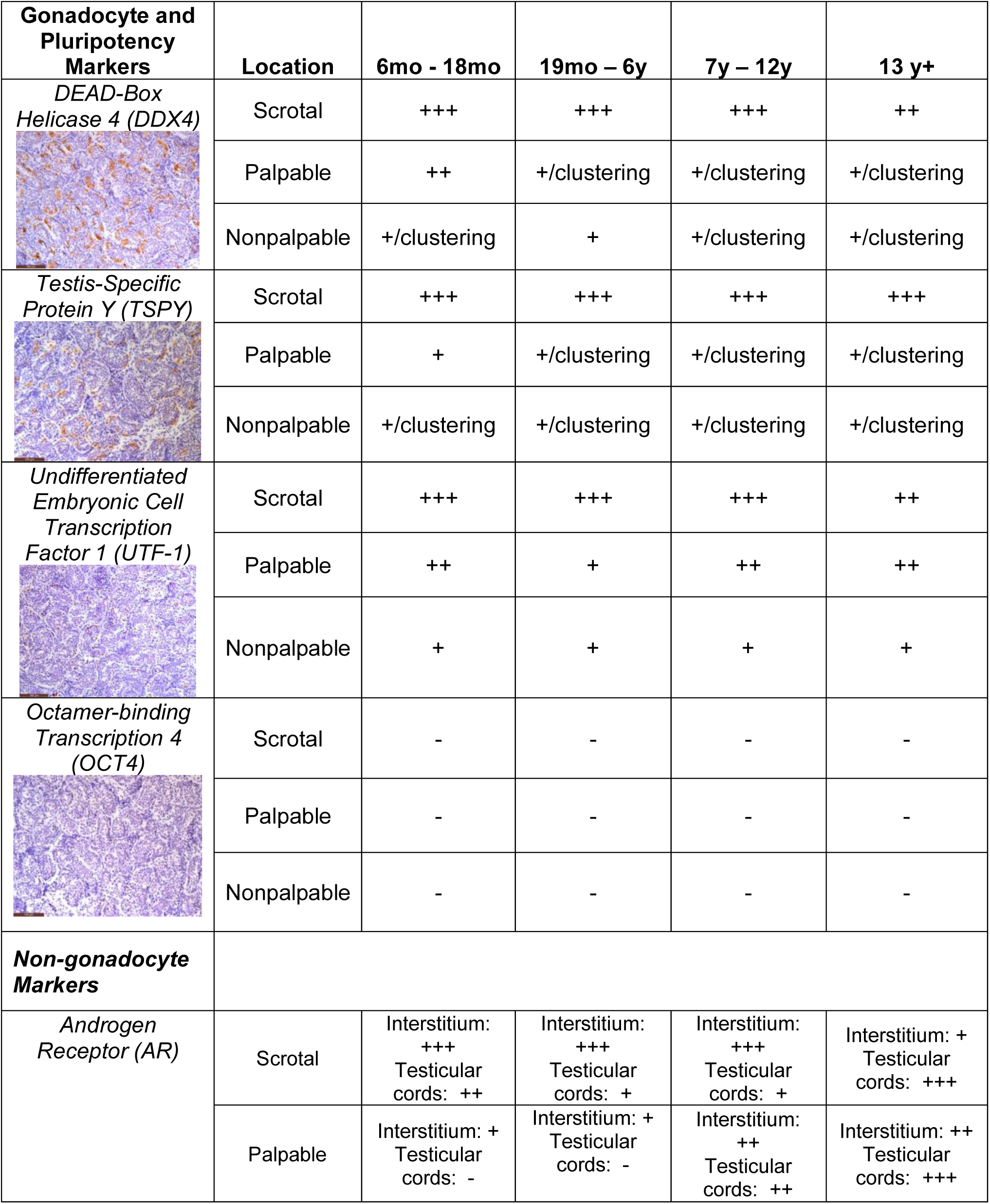
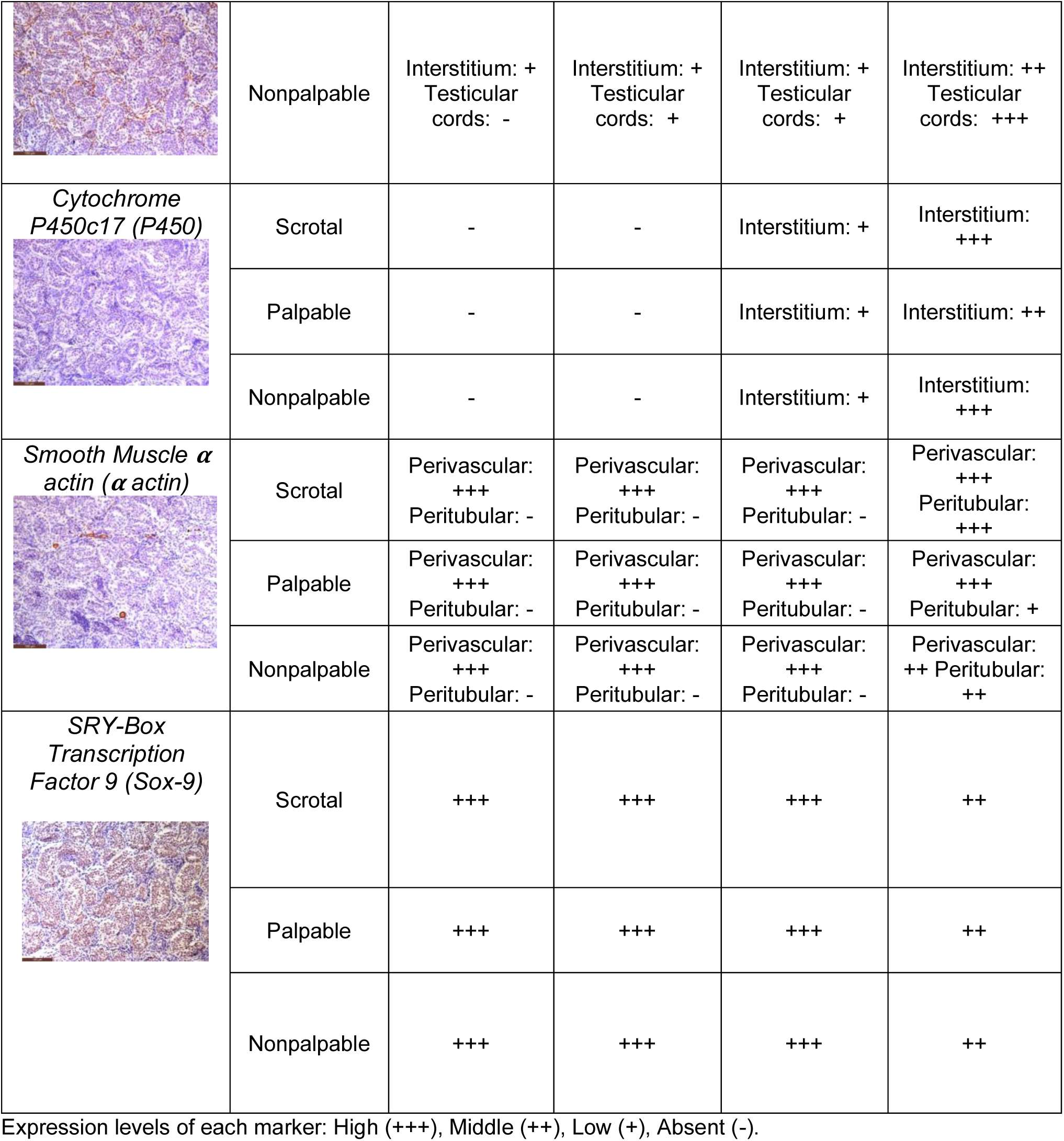
Patterns of Immunohistochemical Staining for Gonadocyte and Nongonadocyte Markers.

|  |  |  |  |  |  |
| --- | --- | --- | --- | --- | --- |
| Gonadocyte and Pluripotency Markers | Location | 6mo - 18mo | 19mo – 6y | 7y – 12y | 13 y+ |
| *DEAD-Box Helicase 4 (DDX4)* | Scrotal | +++ | +++ | +++ | ++ |
|  | Palpable | ++ | +/-clustering | +/-clustering | +/-clustering |
|  | Nonpalpable | +/-clustering | + | +/-clustering | +/-clustering |
| *Testis-Specific Protein Y (TSPY)* | Scrotal | +++ | +++ | +++ | +++ |
|  | Palpable | + | +/-clustering | +/-clustering | +/-clustering |
|  | Nonpalpable | +/-clustering | +/-clustering | +/-clustering | +/-clustering |
| *Undifferentiated Embryonic Cell Transcription Factor 1 (UTF-1)* | Scrotal | +++ | +++ | +++ | ++ |
|  | Palpable | ++ | + | ++ | ++ |
|  | Nonpalpable | + | + | + | + |
| *Octamer-binding Transcription 4 (OCT4)* | Scrotal | - | - | - | - |
|  | Palpable | - | - | - | - |
|  | Nonpalpable | - | - | - | - |
| **Non-gonadocyte Markers** |  |  |  |  |  |
| *Androgen Receptor (AR)* | Scrotal | Interstitium: +++<br>Testicular cords: ++ | Interstitium: +++<br>Testicular cords: + | Interstitium: +++<br>Testicular cords: + | Interstitium: +<br>Testicular cords: +++ |
|  | Palpable | Interstitium: +<br>Testicular cords: - | Interstitium: +<br>Testicular cords: - | Interstitium: ++<br>Testicular cords: ++ | Interstitium: ++<br>Testicular cords: +++ |
|  | Nonpalpable | Interstitial: +<br>Testicular<br>cords: - | Interstitial: +<br>Testicular<br>cords: + | Interstitial: +<br>Testicular<br>cords: + | Interstitial: ++<br>Testicular<br>cords: +++ |
| **Cytochrome**<br>**P450c17 (P450)** | Scrotal | - | - | Interstitial: + | Interstitial:<br>+++ |
|  | Palpable | - | - | Interstitial: + | Interstitial: ++ |
|  | Nonpalpable | - | - | Interstitial: + | Interstitial:<br>+++ |
| **Smooth Muscle $\alpha$**<br>**actin ($\alpha$ actin)** | Scrotal | Perivascular:<br>+++<br>Peritubular: - | Perivascular:<br>+++<br>Peritubular: - | Perivascular:<br>+++<br>Peritubular: - | Perivascular:<br>+++<br>Peritubular:<br>+++ |
|  | Palpable | Perivascular:<br>+++<br>Peritubular: - | Perivascular:<br>+++<br>Peritubular: - | Perivascular:<br>+++<br>Peritubular: - | Perivascular:<br>+++<br>Peritubular: + |
|  | Nonpalpable | Perivascular:<br>+++<br>Peritubular: - | Perivascular:<br>+++<br>Peritubular: - | Perivascular:<br>+++<br>Peritubular: - | Perivascular:<br>++ Peritubular:<br>++ |
| **SRY-Box**<br>**Transcription**<br>**Factor 9 (Sox-9)** | Scrotal | +++ | +++ | +++ | ++ |
|  | Palpable | +++ | +++ | +++ | ++ |
|  | Nonpalpable | +++ | +++ | +++ | ++ |

Expression levels of each marker: High (+++), Middle (++), Low (+), Absent (-).

## Gonadocyte and Pluripotency Markers

Gonadocyte markers, DDX4 and TSPY, shared similar patterns and location of expression across all specimens. At ages 12 and younger, the germ cells stained were part of the architectural structure within the testicular cords. For specimens in ages 13 or older, the start of puberty is evidenced by the development of an increasingly wider lumen of seminiferous tubules. Expression patterns noted in detail in **Supplemental Figure 1-2.** Unlike DDX4 and TSPY, germ cell marker OCT-4 had negative staining in both scrotal and UDT.

### DEAD-box helicase 4 (DDX4)

The scrotal testicular specimens at all ages had a greater number of germ cells staining positive for DDX4 and localized to the lumen and basal lamina of all testicular cords (**Supplemental Fig 1. A, D, G, J).** At ages 6-18months, palpable UDT had significant interstitial fibrosis with preserved staining of DDX4 positive cells, whereas nonpalpable testis had decreased staining (**Supplemental Fig 1. B-C**). At older ages after 18 months, UDT showed increased stromal fibrosis with more severe reduction of germ cell expression relative to scrotal tissue (**Supplemental Fig 1. D-L**). At ages 13 or older, the scrotal testis was characterized by a wide lumen corresponding to the development of seminiferous tubules, and continued expression of DDX4 confined within the tubules (**Supplemental Fig 1. J).** In contrast, palpable and nonpalpable UDT had fewer seminiferous tubules with positive DDX4 positive germs cell (**Supplemental Fig 1. K, L**). In specific specimens, the DDX4 positive cells were exclusively localized to specific tubules in a cluster spatial arrangement. (**Supplemental Fig 1. E, H, K**). This finding was seen in 48 % (15/31) of UDT specimens (**Fig 1. A-D; Supplemental Table 3**), and it was not associated with age or testicular location.

**Figure 1.**
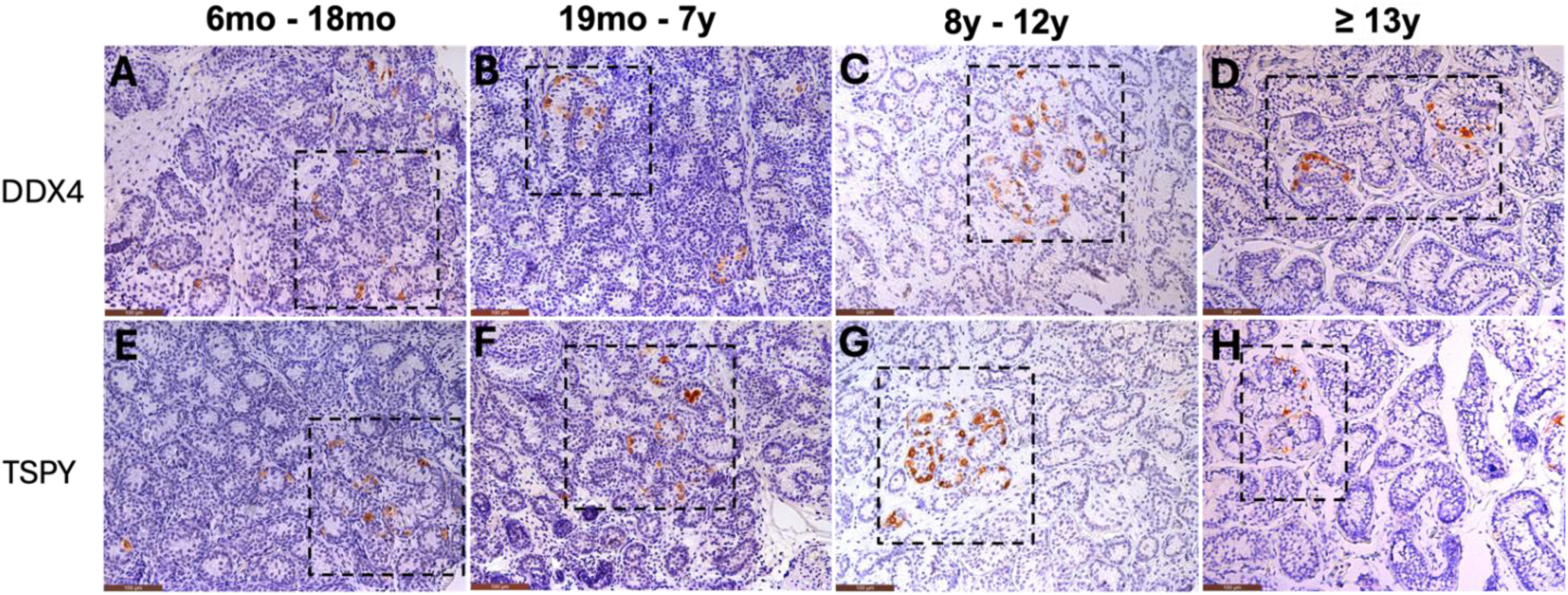
Gonadocyte Clustering in Cryptorchid Testis. Cryptorchid testicular specimens shown in A-B exhibit DDX4 expression, and E-H exhibit TSPY expression. DDX4 and TSPY stain positive for germ cells, highly specialized cells that sustain gamete production and spermatogenesis. DDX4, also known as VASA protein, is highly specific and reliable marker for germ cells in human testis. TSPY is a male specific germ cell marker that serves important functions in gonocyte and spermatogonia differentiation. Square boxes highlight the germ cell clusters, a common finding seen in 15 out of 31 (48%) specimens with this pathology. These images were reproduced at 20x magnification. Apart from the focal regions containing positively stained germ cell clusters, the remaining seminiferous tubules in the tissue section were devoid of germ cells. No scrotal testes had this finding of germ cell clustering on staining (no shown).

### Testis-Specific Protein Y (TSPY)

TSPY, male-specific germ cell marker, expression was similar to that of DDX4. Across all ages and independent of location, UDT had decreased levels of TSPY germ cell expression (**Supplemental Fig 2. A-L**). In comparison to DDX4, fewer germ cells were stained for this marker, and when expressed these cells appeared to be confined to the basal lamina of the seminiferous tubules (**Supplemental Fig 2. A, D, G, J).** Palpable and non-palpable UDT showed a variable level of stromal fibrosis across all ages that was more evident in older nonpalpable testis at ages 19 months to 7 years and 8 years to 12 years (**Supplemental Fig 2. A-I**). As seen with DDX4, clustering of TSPY cells were seen in 48% of UDT specimens (**Fig 1. E-H, Supplemental Table 3**).

### Undifferentiated Embryonic Cell Transcription Factor 1 (UTF-1)

At 12 months, UTF-1was expressed in the nuclei of certain germ cells located in the basal lamina of the testicular cords, as indicated by co-expression with cytoplasmic germ cell marker DDX4 in scrotal testis (**Supplemental Fig. 3).** While the levels of UTF-1 expression varied across scrotal specimens and appeared to decrease with age, all testicular cords contained germ cells that stained positive for UTF-1. (**Fig 2. A, D, G, J**). At 6-18 months, UTF-1 expression was primarily identified in the basal lamina in the periphery of testicular cords (**Fig 2. A**). From ages 19 months-12 years, UTF-1 expression was found throughout the cords (**Fig 2. D, G**). By age 13 years, the formation of the lumen occurs, and the UTF-1 nuclei return to initial location in the basal lamina (**Fig 2. J**). UTF-1 expression in UDT was either reduced or fully absent, and these findings were seen at all ages and whether the testis was palpable or nonpalpable. (**Fig 2. A-L**)

**Figure 2.**
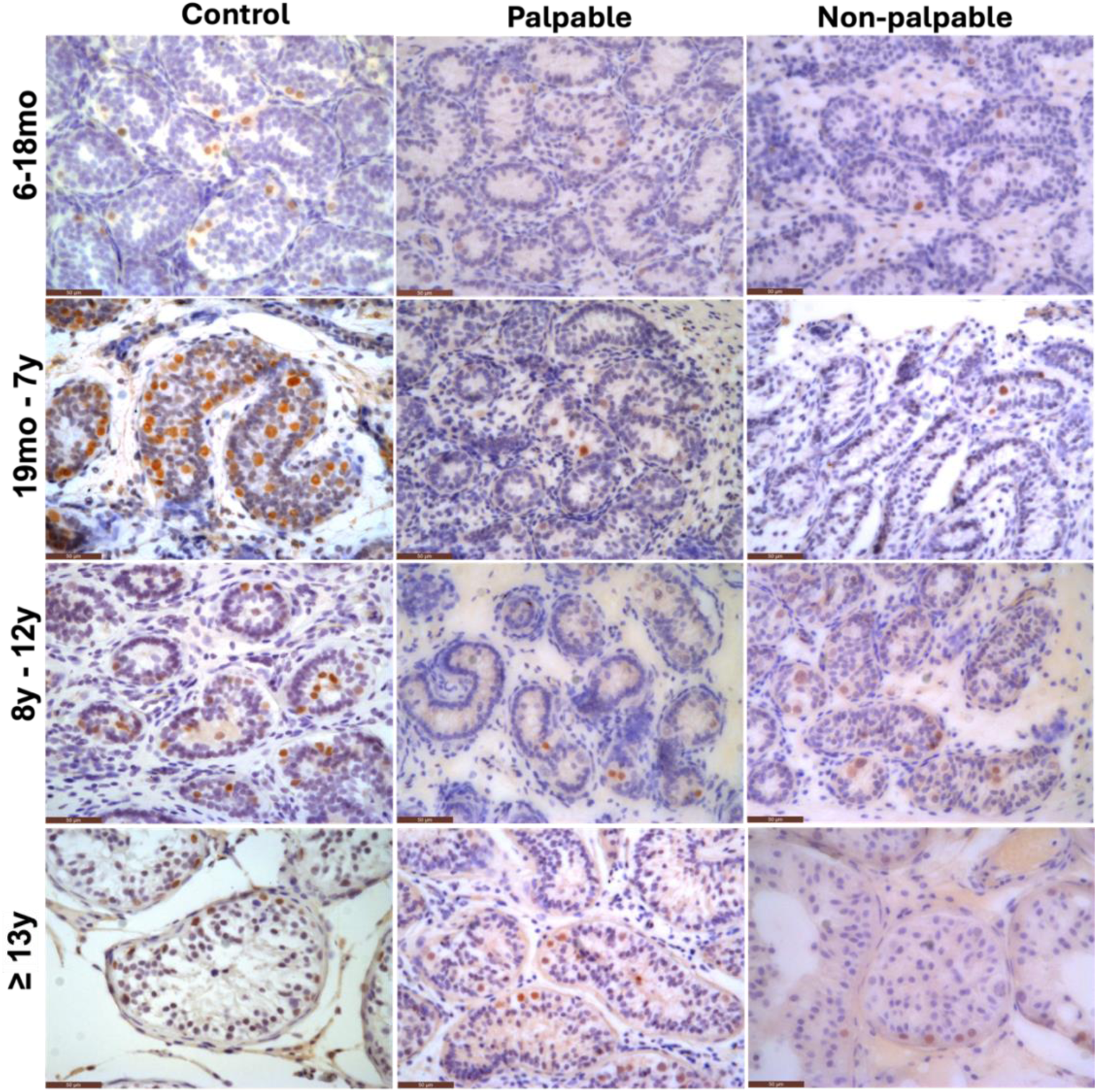
UTF-1 Expression Patterns in Cryptorchid Testis by Age and Location. Cryptorchid specimens were stained for pluripotency marker UTF-1. UTF-1 is a transcriptional regulator critical for early gonadal development. In mammals, UTF-1-positive spermatogonia are expected to be present prepubertally, and nearly absent after puberty. In our control group of scrotal testis, UTF-1 maintained moderate of expression within testicular cords at all ages, particularly between 6-months to 7 years. In cryptorchid testis, TSPY expression was reduced across all following ages at time of surgery: 6-18 months, 19 months – 7 years, 8-12 years, and 13 years or older. Images are shown at 40x magnification.

### Age and Location effect on Germ Cell Count

Mean germ cell per tubule varied by age group and testicular location (scrotal, palpable UDT, and nonpalpable UDT). In infants aged 6 to 18 months, mean germ cell counts per tubule expression of DDX4, TSPY, and UTF-1 were relatively similar across testicular locations (**Table 2**). Across all other age groups, scrotal testes consistently exhibited higher germ cell counts per tubule, as indicated by DDX4, TSPY, and UTF-1 expression, compared to both palpable and nonpalpable undescended testes (UDT) (**Table 2)**.

**Table 2.** Comparison of mean number of germ cell per tubule by testicular location.

| **Age Groups** | **Scrotal** | **Palpable** | **Nonpalpable** |
| --- | --- | --- | --- |
| *6 months - 18 months* |  |  |  |
| DDX4 | 3.2 | 3.27 | 3.77 |
| TSPY | 2.95 | 3.73 | 3.53 |
| UTF-1 | 2.55 | 1.97 | 1.47 |
| *19 months - 6 years* |  |  |  |
| DDX4 | 6.6 | 2.3 | 0.9 |
| TSPY | 8.4 | 1.67 | 1.3 |
| UTF-1 | 12.5 | 1.2 | 0.3 |
| *7 years – 12 years* |  |  |  |
| DDX4 | 6.1 | 3.06 | 2.12 |
| TSPY | 5.1 | 2.94 | 2.23 |
| UTF-1 | 4 | 0.84 | 0.7 |
| *13 years +* |  |  |  |
| DDX4 | 8.9 | 3.37 | 4.42 |
| TSPY | 8.2 | 2.17 | 4.1 |
| UTF-1 | 1.8 | 2.27 | 0.7 |

<sup>a</sup> Statistical significance was not able to be determined due to small sample size of control (n=1 per age group).

## Non-gonadocyte Markers

The characterization of the somatic architecture of UDT was facilitated by the expression patterns of SOX9, AR, α-SMA, CYP17A1. Androgen Receptor (AR) expression was overall diminished in UDT relative to scrotal testis. However, both shared similar localizations as scrotal testis, exhibiting rising levels of expression within tubular cords at older ages, particularly in the peripubertal ages (+13yo) (**Table 2, Supplemental Figure 4**). There were no qualitative differences in expression of SOX9, α-SMA, and CYP17A1 (**Table 2, Supplemental Figure 5 – 8**).

## Discussion

This study maps spatial and temporal expression patterns of germ cell, pluripotency, and somatic markers in postnatal cryptorchid testes, directly compared to age-matched scrotal controls. Unlike prior studies that examined limited markers, we included a comprehensive panel spanning germ cell identity (DDX4, TSPY), pluripotency (UTF-1), and somatic niche components (SOX9, AR, α-SMA, CYP17A1). This panel builds on our previously published normal testicular development framework and enables a more comprehensive view of postnatal testicular maturation and dysgenesis in the setting of cryptorchidism ^15^. Collectively, our findings support a model in which cryptorchidism involves both germ cell depletion and disrupted SSC lineage formation, with UTF-1 downregulation and germ cell clustering as early signatures of testicular dysgenesis.

### Germ Cell Dysgenesis in Cryptorchidism

Germ cell loss is a hallmark feature of UDT pathology, and our findings support a model of progressive, spatially heterogeneous dysgenesis during postnatal development.^9^ We observed a decline in expression of DDX4 and TSPY in cryptorchid testes that correlated with both age and anatomical location, with the greatest reductions seen in nonpalpable (presumably intra-abdominal) testes beyond 18 months of age (**Table 1, Supplemental Figure 1**). In our cohort, cryptorchid specimens exhibited both quantitative loss of signal and uneven distribution relative to scrotal testis, suggesting that germ cell loss in cryptorchidism may not occur uniformly across the seminiferous epithelium.

A novel observation in this study was the presence of germ cell clustering, where DDX4– and TSPY-positive cells were restricted to a minority of seminiferous cords while the majority were entirely devoid of germ cells (**Supplemental Figure 2**). This pattern was absent in all scrotal controls. These clusters may represent focal preservation of the germ cell lineage in protected microenvironments or regions of delayed degeneration. Similarly, Ferragut Cardoso et al. (2021) described mixed histologic patterns in undescended testes, with some seminiferous tubules displaying Sertoli-only histology and others retaining partial spermatogenesis, reinforcing the concept of localized survival ^17^. We speculate that this phenotype may arise from disrupted somatic support, particularly alterations in Sertoli-germ cell signaling, microenvironments, or regional thermal gradients. Histologic and transcriptomic evidence from human and experimental models suggest that Sertoli cell immaturity, niche fibrosis, and abnormal paracrine signaling may contribute to focal germ cell attrition or survival ^18–20^.

Mechanistically, our findings align with models in which elevated intra-abdominal temperature and disrupted paracrine cues impair germ cell survival through non-apoptotic pathways. Experimental studies have shown that heat stress leads to germ cell cell-cycle arrest, defects in spatial arrangement of testicular cells, impaired DNA synthesis, and meiotic disruption ^21–23^. The focal clustering we observed may therefore mark regions where germ cells temporarily evade these stressors or where somatic support remains partially intact. Focal spermatogenesis in humans has only been previously reported in topographical data from patients with Klinefelter syndrome, and provides a rationale for high success rates in testicular sperm extraction in these patients ^24^. Clinically, early germ cell clustering and regional depletion may serve as histological biomarkers of early testicular dysgenesis, particularly in children undergoing biopsy or fertility preservation. If validated, this pattern could help identify testes at highest risk for germ cell loss and support earlier orchiopexy or targeted SSC-preserving therapies.

### UTF-1 and Pluripotency Marker Dynamics

UTF-1 is a chromatin-associated transcription factor expressed in pluripotent cells^25^ and a subset of undifferentiated spermatogonia during early testicular development. In the rat testis, UTF-1 is highly expressed in neonatal gonocytes and becomes progressively restricted to early spermatogonia during postnatal maturation, particularly those co-expressing ZBTB16 and retaining self-renewal potential. ^26^ This population includes As, Apr, and short Aal spermatogonia, the foundational units of the spermatogonial stem cell (SSC) pool. Similar developmental restriction has been documented in humans, and the reduced expression of this spermatogonial stem cell marker has been associated with impaired spermatogenesis.^13,16,27^ More recently, an adult human transcriptional cell atlas using single cell RNA sequencing demonstrates that UTF-1 is highly expressed in a novel and early quiescent state of SSC, ^28^ suggesting this marker plays a pivotal role in the development and maturation of spermatogonia.

In our study, UTF-1 expression was consistently strong in scrotal testes across all age groups, whereas both palpable and nonpalpable undescended testes (UDT) showed significantly reduced or absent UTF-1 staining, especially in patients older than 18 months (**Figure 2 B–C**). This pattern mirrors findings in rodent models, where UTF-1 expression is rapidly downregulated beyond the neonatal period and is absent from differentiating spermatogonia and spermatocytes ^26^. In our age-stratified analysis, UTF-1-positive cells were virtually absent from UDT after early infancy, while persisting in age-matched scrotal controls (**Table 1**) suggesting that loss of UTF-1 may reflect failure of the gonocyte-to-SSC transition under dysregulated developmental conditions.

Functionally, UTF-1 plays a role in maintaining an epigenetic landscape permissive to pluripotency. Studies in embryonic stem cells and early germ cells indicate that UTF-1 interacts with chromatin remodeling complexes and modulates nucleosome compaction, thereby maintaining transcriptional plasticity rather than directly regulating proliferation ^29^. In this context, the loss of UTF-1 in UDT may reflect failure to initiate or sustain the chromatin configuration necessary for SSC identity. This interpretation is consistent with observations that UTF-1 is downregulated even in morphologically intact germ cells in non-scrotal testes, indicating functional exhaustion or altered cell fate potential ^29,30^. Loss of UTF-1 may mark both the absence of undifferentiated germ cells and failure of epigenetic programming essential for SSC establishment. Thus, UTF-1 downregulation may serve as an early molecular signature of germ cell dysgenesis in cryptorchidism.

### Somatic Cell Marker Expression and Niche Disruption

Sex determination and early testis differentiation are driven by SRY-mediated upregulation of SOX9, which is essential for Sertoli cell specification and the organization of testicular cords ^31,32^. SOX9 also promotes differentiation of Leydig cells and interstitial structures ^33^. These early events create the foundation for the testicular niche. While SOX9-positive Sertoli cells were present in all samples, its expression appeared disorganized in UDT, particularly in nonpalpable specimens. This morphological alteration parallels experimental findings in which Sertoli cells lose apical-basal polarity and junctional integrity under cryptorchid conditions ^34^.

Leydig cell function was assessed via CYP17A1 and AR staining. CYP17A1, typically active during mini-puberty and puberty, was absent in all samples from 6 months to 7 years and reappeared in adolescence, consistent with its expected developmental pattern ^15^. Expression in older UDT was reduced compared to scrotal controls, suggesting impaired reactivation of steroidogenesis.

AR was strongly expressed in interstitial cells of scrotal testes and showed variable expression in Sertoli cells depending on age. In UDT, AR expression was reduced in both compartments, especially in nonpalpable specimens. Given the role of AR in maintaining Sertoli function and germ cell adhesion, diminished signaling may contribute to broader niche instability.

Peritubular myoid cells were assessed using α-SMA. In prepubertal samples, α-SMA staining was restricted to vascular smooth muscle. In scrotal testes older than 13 years, peritubular α-SMA became evident, consistent with post-pubertal activation of PMC function ^15^. This pattern was reduced or absent in age-matched UDT, suggesting delayed PMC maturation. Rodent transcriptomic data support this, showing downregulation of smooth muscle markers and impaired ECM remodeling in cryptorchidism ^34^.

Collectively, these findings suggest somatic niche dysfunction in cryptorchidism, potentially contributing to UTF-1 loss, disrupted Sertoli cell structure, impaired Leydig and PMC signaling, and localized germ cell clustering.

### Limitations

Several limitations should be noted. First, sample sizes were small within age groups, especially for older scrotal controls, limiting statistical power and generalizability. Second, the cross-sectional design prevents assessment of longitudinal changes of marker expression and differentiation between delayed maturation or irreversible germ cell loss.

Third, while IHC provides spatial resolution, it is inherently semi-quantitative and influenced by antibody sensitivity and tissue preservation, particularly affecting detection of temporally restricted markers like CYP17A1 and UTF-1. Fourth, we did not include co-staining with additional SSC markers (e.g., SALL4, GFRA1, ID4) or perform transcript-level validation. As such, we cannot fully resolve the functional identity of UTF-1-negative germ cells.

Fifth, some scrotal controls had congenital abnormalities or unrelated pathology, which may have influenced histologic profiles. Finally, Tanner stage data were unavailable, which limits our ability to align pubertal status with marker expression in the older age groups.

Despite these limitations, the consistent findings of germ cell clustering and UTF-1 loss highlight early hallmarks of testicular dysgenesis in cryptorchidism. These features may help identify high-risk patients and provide a rationale for earlier orchiopexy or SSC-preserving strategies, while guiding future mechanistic and translational studies.

## Author’s Roles

P.A.S, M.M, M.C., C.D, J.F.S, L.S.B, Y.L contributed to study design and development, acquisition of data, analysis and interpretation of data, manuscript drafting and final approval of the version to be published; P.A.S executed the study.

## Acknowledgements

The authors gratefully acknowledge the UCSF Department of Urology for their generous support and access to departmental resources that made this study possible. We also thank Dr. Gerald Cunha for his thoughtful discussions on male gonadal development and for offering valuable insights that helped shape the future directions of this research.

## Abbreviations

A-actin: Alpha-actin
AR: Androgen Receptor
DDX4: DEAD-box helicase 4
IHC: Immunohistochemistry
OCT4: Octamer-binding transcription factor 4
P450: Cytochrome P450
SSC: Spermatogonial Stem Cells
SOX9: SRY-box transcription factor 9
TSPY: Testis-Specific Protein Y-encoded
UDT: Undescended Testes
UTF-1: Undifferentiated Embryonic Cell Transcription Factor 1

