## Supplemental Table 1 for "“Mapping of Gonadal Development in Cryptorchidism: UTF-1 and Germ Cell Dysgenesis”"

**Supplementary Table 1. Testis Specimens used in this study by age and location**

| Specimen Number | Age at time of Biopsy | Location |
|---|---|---|
| ***Scrotal Testis (Control)*** | | |
| S1 | 12 mo | scrotal |
| S2 | 16 mo | scrotal |
| S3 | 6.5 y | scrotal |
| S4 | 11 y | scrotal |
| S5 | 13 y | scrotal |
| ***Cryptorchid Testis*** | | |
| *Optimal Surgical Window (6-18 months)* | | |
| UDT1 | 8 mo | non-palpable |
| UDT2 | 11 mo | palpable |
| UDT3 | 12 mo | palpable |
| UDT4 | 12 mo | non-palpable |
| UDT5 | 13 mo | palpable |
| UDT6 | 18 mo | non-palpable |
| *Pre-Puberty (19 months- 7 years)* | | |
| UDT7 | 1.6 y | palpable |
| UDT8 | 2 y | palpable |
| UDT9 | 2 y | non-palpable |
| UDT10 | 2.5 y | non-palpable |
| UDT11 | 3 y | palpable |
| UDT12 | 3 y | palpable |
| UDT13 | 4 y | palpable |
| UDT14 | 4 y | palpable |
| UDT15 | 7 y | non-palpable |
| *Peri-Puberty (8years - 12 years)* | | |
| UDT16 | 8 y | palpable |
| UDT17 | 8 y | non-palpable |
| UDT18 | 8 y | non-palpable |
| UDT19 | 8 y | non-palpable |
| UDT20 | 10 y | palpable |
| UDT21 | 10 y | palpable |
| UDT22 | 10 y | palpable |
| UDT23 | 11 y | palpable |
| UDT24 | 12 y | palpable |
| UDT25 | 12 y | palpable |
| *Post-Puberty (≥13 years)* | | |
| UDT26 | 13 y | non-palpable |

| | | |
|---|---|---|
| UDT27 | 14 y | palpable |
| UDT28 | 14 y | palpable |
| UDT29 | 16 y | non-palpable |
| UDT30 | 16 y | non-palpable |
| UDT31 | 18 y | palpable |
