## Supplemental Table 2 for "“Mapping of Gonadal Development in Cryptorchidism: UTF-1 and Germ Cell Dysgenesis”"

**Supplementary Table 2. Antibodies used in the Immunohistochemical Mapping of Cryptorchidism**

| Antibody Name | 6/24/2026 3:19:00 PMConcentration | Catalog No. | Company |
|---|---|---|---|
| **Gonadocyte and Pluripotency Markers** | | | |
| DEAD-Box Helicase 4 | 1:500 | ab13840 | Abcam |
| Testis-Specific Protein Y | 1:2000 | NA | Lau Lab, UCSF |
| Undifferentiated Embryonic Cell Transcription Factor 1 | 1:100 | AF3958 | R&D Systems |
| **Nongonadocyte Markers** | | | |
| Androgen Receptor | 1:400 | sc-816 | Santa Cruz Biotechnology |
| Cytochrome P450c17 | 1:150 | NBP2-01151 | Novus Biologicals |
| Octamer--binding Transcription 4 | 1:100 | sc-5279 | Santa Cruz Biotechnology |
| Smooth Muscle α actin | 1:2000 | A5228-200UL | Sigma |
| SRY-Box Transcription Factor 9 | 1:1000 | AF3075 | R&D Systems |
| Biotinylated Donkey anti-Rabbit | 1:100 | RPN1004V | GE Healthcare UK Limited |
| Biotinylated Sheep anti-Mouse | 1:100 | RPN1001V | GE Healthcare UK Limited |
| Biotinylated Donkey anti-Goat | 1:100 | sc-2042 | Santa Cruz Biotechnology |

a. Dr. Chris Lau, UCSF, Department of Medicine, San Francisco, CA (Lau et al., 2000).
