## Supplemental Figure 3 for "“Mapping of Gonadal Development in Cryptorchidism: UTF-1 and Germ Cell Dysgenesis”"

**Supplementary Figure 3. Co-expression of UTF-1 and DDX4 in control testicular specimen**

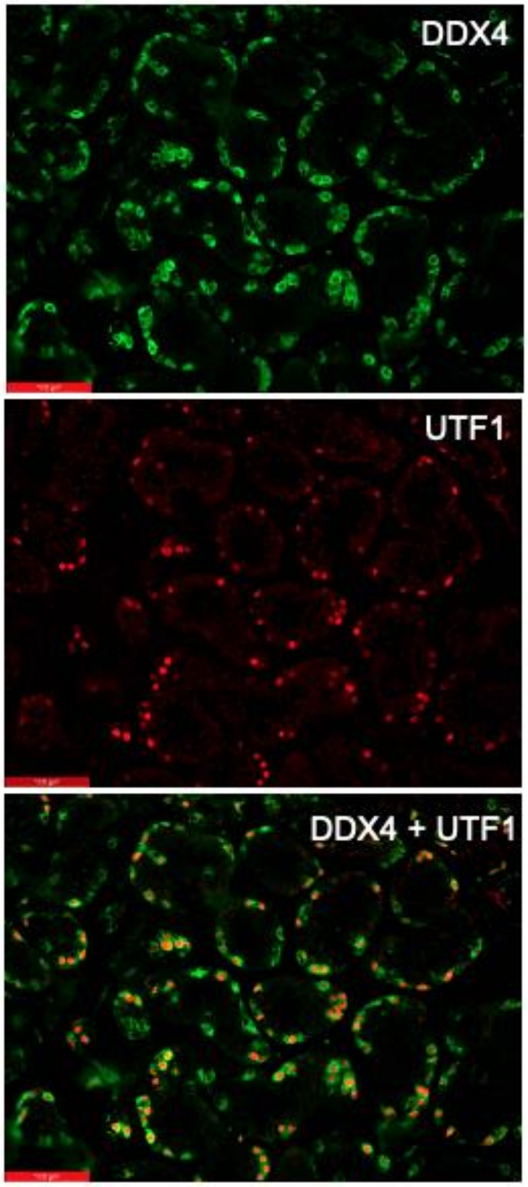


Immunofluorescence staining of a 12-month-old control testis shows co-localization of UTF-1 (red) and DDX4 (green) in germ cells. Merged images reveal overlapping expression of UTF-1 and DDX4 in a subset of cells, indicating the presence of undifferentiated germ cells. Scale bar: 40x magnification.
