## Supplemental Figure 4 for "“Mapping of Gonadal Development in Cryptorchidism: UTF-1 and Germ Cell Dysgenesis”"

**Supplementary Figure 4. Expression of AR in Cryptorchid Testis by Age and Location**

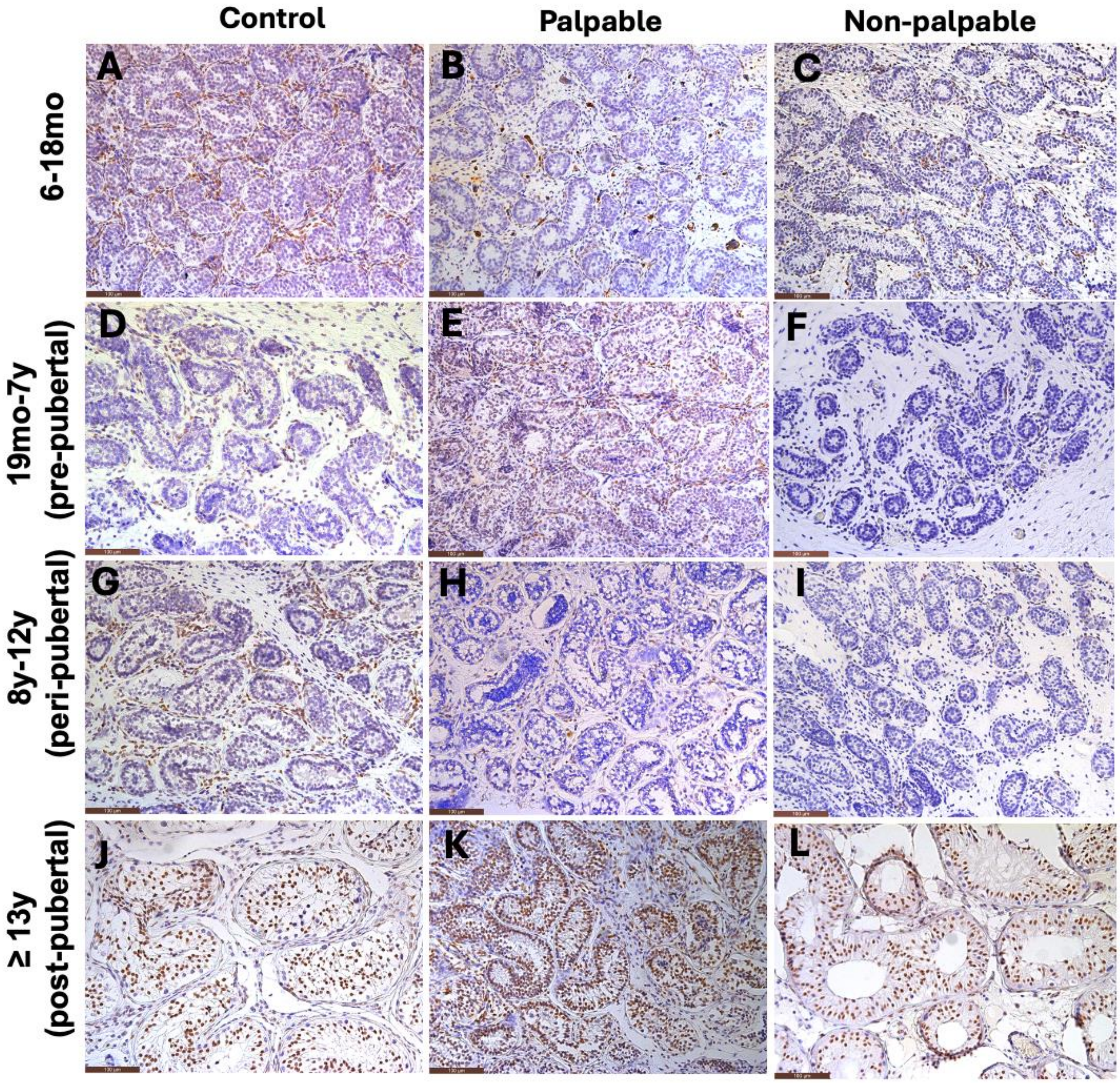


Cryptorchid specimens were stained for androgen receptor (AR). In scrotal testis, AR was expressed in the interstitium and testicular cords across all ages. At age 13, AR was predominantly expressed in the testicular cords. In comparison to scrotal testis, both palpable and non-palpable cryptorchid testis showed reduced AR expression at ages 18 months (A-C), 19 months -7 years (D-F), and 8-12 years (G-I), and 13 years or older (J-L). Testicular cord expression of AR was similar between scrotal and cryptorchid testis at ages 13 years or older (J-L).

**Supplementary Figure 5. Expression of α-SMA in Cryptorchid Testis by Age and Location**

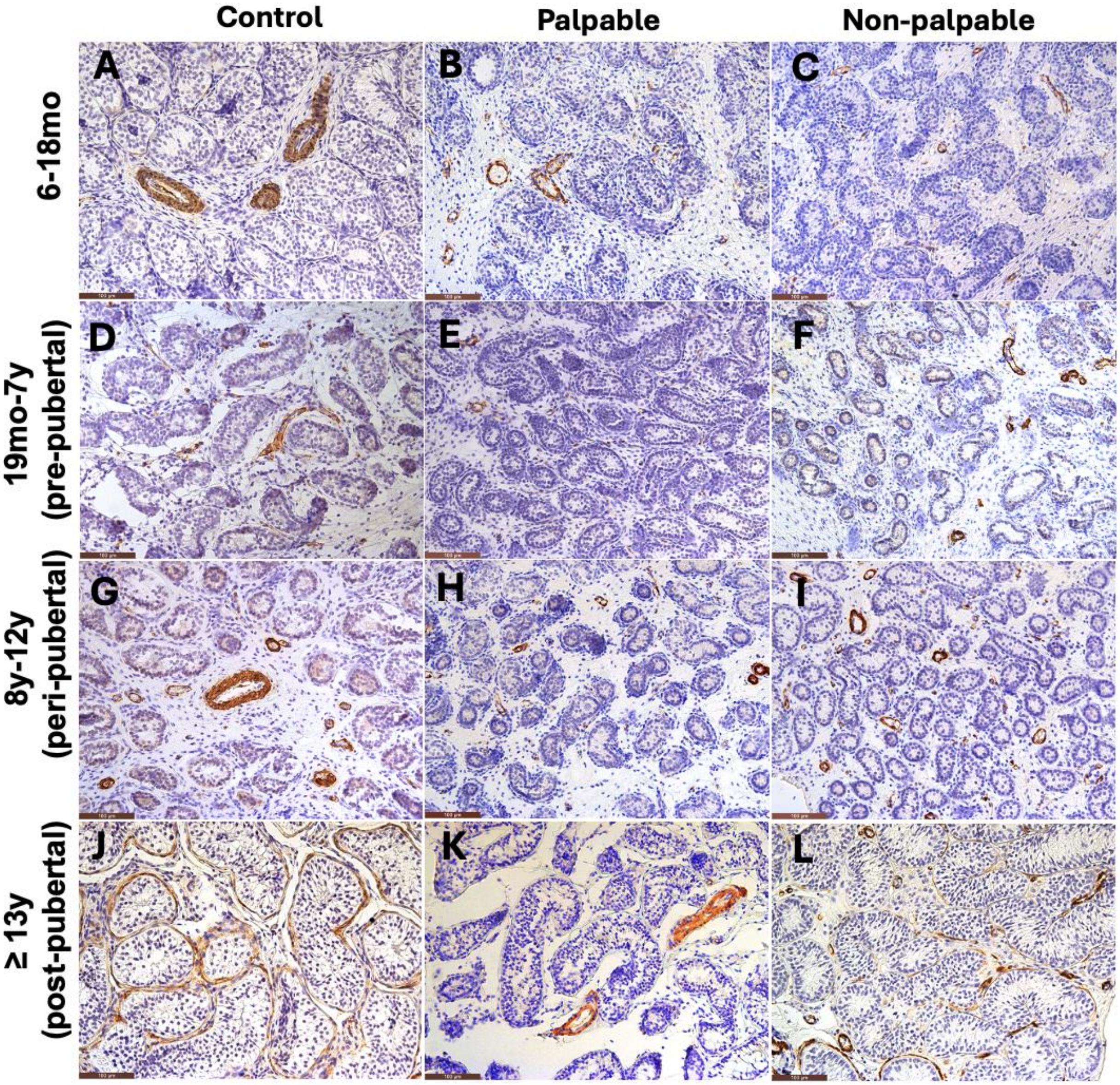


Cryptorchid specimens were stained for non-gonadocyte marker, *Smooth Muscle* ***α*** *actin (****α*** *actin*). Scrotal testis and cryptorchid testis showed positive perivascular staining within testicular cords at the following ages: 6-18 months (A-C), 19 months -7 years (D-F), 8-12 years (G-I), and 13 years or older (J-L). Peritubular expression of ***α****-actin was seen only at ages* 13 years or older (J-L).

**Supplementary Figure 6. Expression of CYP17A1 in Cryptorchid Testis by Age and Location**

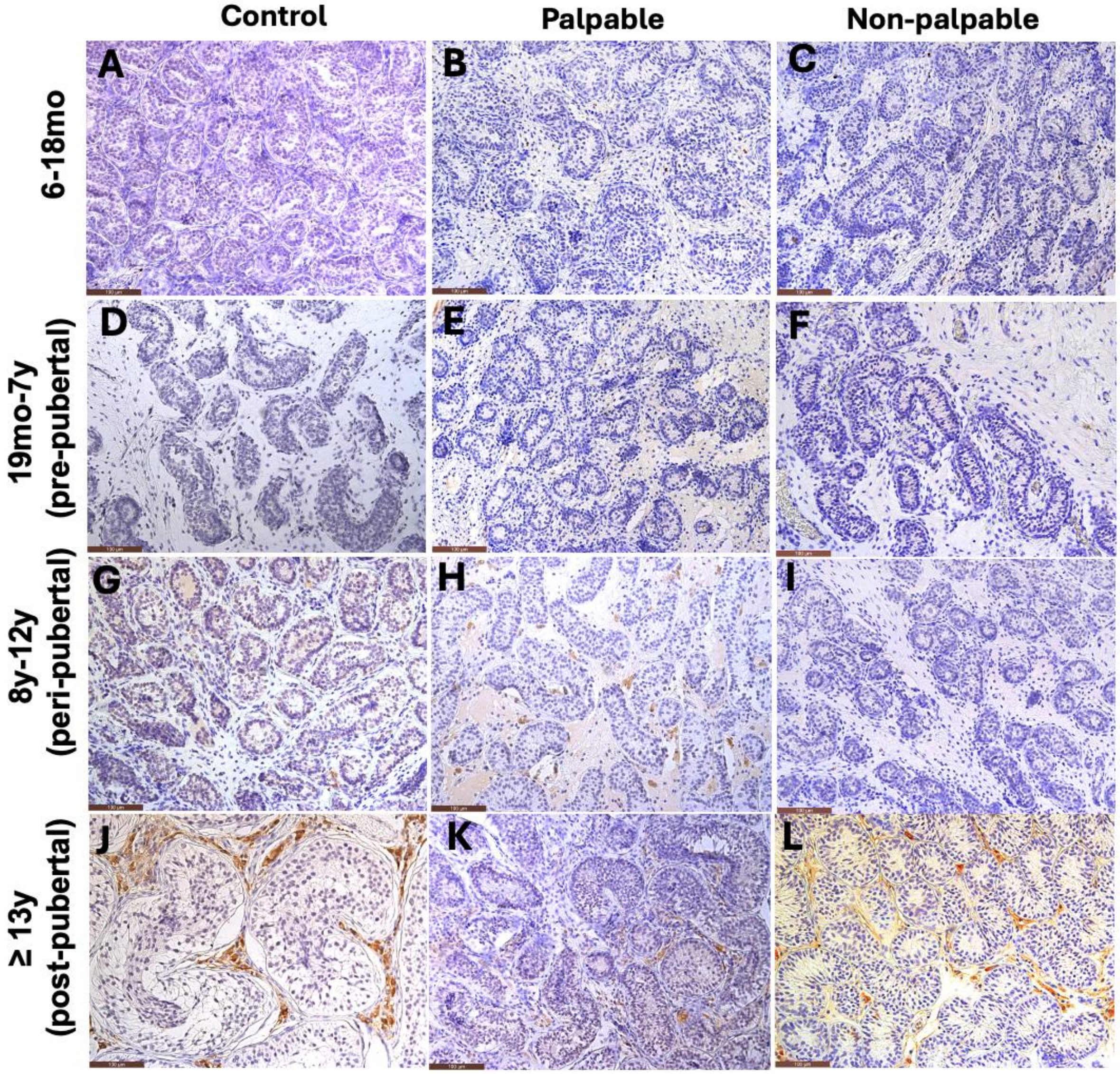


Cryptorchid specimens were stained for *Cytochrome P450c17 (P450).* CYP17A1 expression was negative in the younger age groups. At ages 8-12 years (G-I) and 13 years or older (J-L), there was a moderate increase in expression of CYP17A1 in the interstitium. These patterns were consistent in scrotal and cryptorchids testis.

**Supplementary Figure 7. Expression of SOX-9 in Cryptorchid Testis by Age and Location**

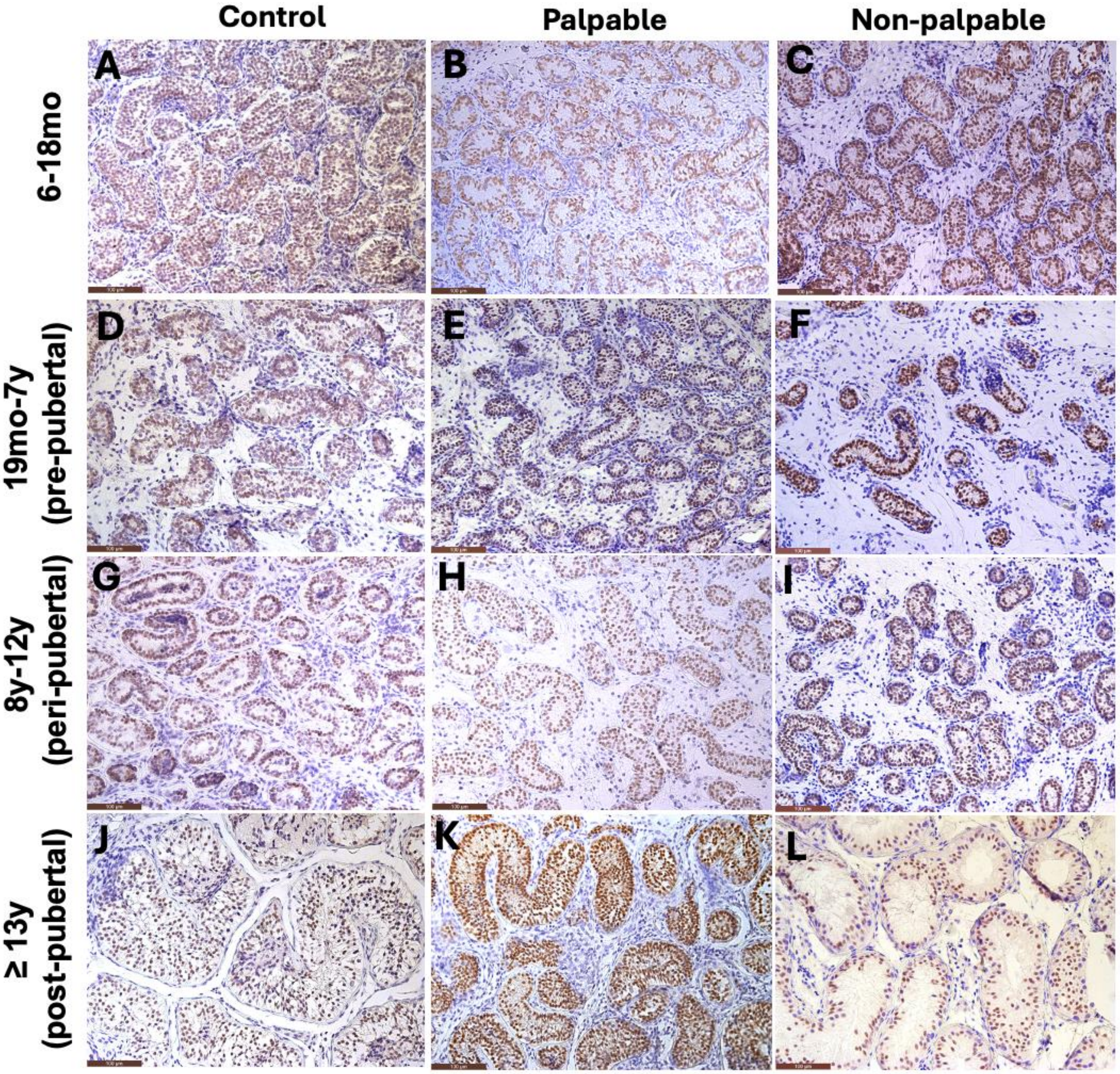


Cryptorchid specimens were stained for *SRY-Box Transcription Factor 9 (Sox-9).* Scrotal testis and cryptorchid testis showed similar levels of SOX-9 expression within testicular cords at all ages: 6-18 months (A-C), 19 months -7 years (D-F), 8-12 years (G-I), and 13 years or older (J-L).

**Supplementary Figure 8. Expression of OCT4 in Cryptorchid Testis by Age and Location**

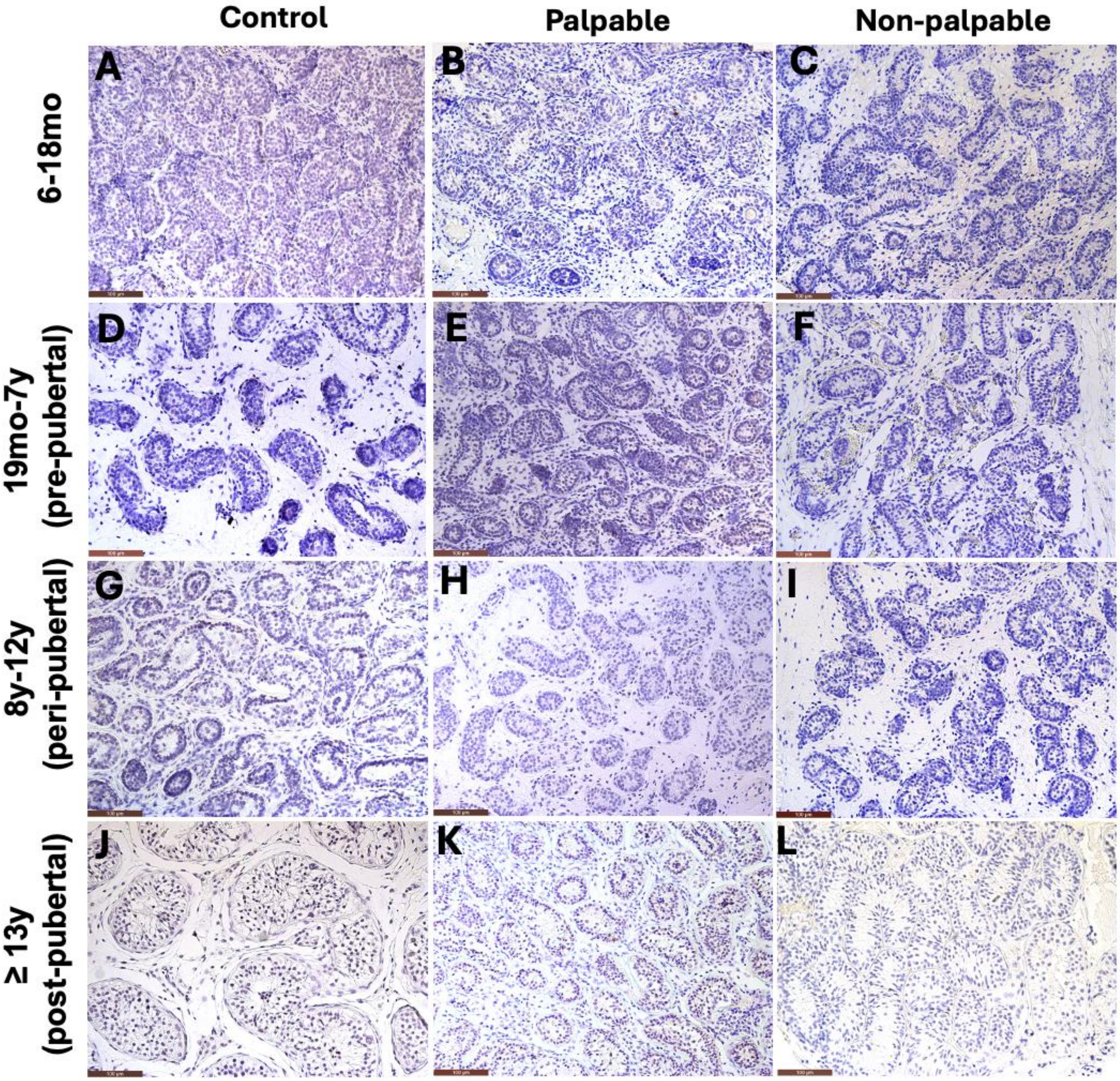


Cryptorchid specimens were stained for *Octamer-binding Transcription 4 (OCT4)*. No expression of OCT 4 marker was seen in scrotal and cryptorchid testis at all ages: 6-18 months (A-C), 19 months -7 years (D-F), 8-12 years (G-I), and 13 years or older (J-L).
